# Lifespan differences in visual short-term memory load-modulated functional connectivity

**DOI:** 10.1101/2022.08.31.506084

**Authors:** Selma Lugtmeijer, Linda Geerligs, Kamen A. Tsvetanov, Daniel J. Mitchell, Cam-CAN, Karen L. Campbell

**Author notes:** Corresponding author: Selma Lugtmeijer, Department of Psychology, Brock University, 1812 Sir Isaac Brock Way, St. Catharines, ON L2S 3A1, Canada.

## Abstract

Working memory is critical to higher-order executive processes and declines throughout the adult lifespan. However, our understanding of the neural mechanisms underlying this decline is limited. Recent work suggests that functional connectivity between frontal control and posterior visual regions may be critical, but examinations of age differences therein have been limited to a small set of brain regions and extreme group designs (i.e., comparing young and older adults). In this study, we build on previous research by using a lifespan cohort and a whole-brain approach to investigate working memory load-modulated functional connectivity in relation to age and performance. The article reports on analysis of the Cambridge Centre for Ageing and Neuroscience (Cam-CAN) data. Participants from a population-based lifespan cohort (*N*=111, age 23-86) performed a visual short-term memory task during functional magnetic resonance imaging. Visual short-term memory was measured with a delayed recall task for visual motion with three different loads. Whole-brain load-modulated connectivity was estimated using psychophysiological interactions in a hundred regions of interest, sorted into seven networks (Schaefer et al., 2018, Yeo et al., 2011). Results showed that load-modulated functional connectivity was strongest within the dorsal attention network followed by the visual network during encoding and maintenance. With increasing age, load-modulated functional connectivity strength decreased throughout the cortex. Within the dorsal attention network, increased load-modulated connectivity strength was related to better task performance in an age-invariant way. Our results demonstrate the widespread negative impact of age on the modulation of functional connectivity by working memory load. Older adults might already be close to ceiling in terms of their resources at the lowest load and therefore less able to further increase connectivity with increasing task demands.

**Highlights:**

- We examine visual short-term memory load-related functional connectivity
- And how age and performance affect modulation of connectivity by memory load
- Modulation of connectivity is strongest in dorsal attention and visual networks
- Load-modulated connectivity strength decreases with increasing age
- Load-modulated connectivity in the dorsal attention network relates to performance

## 1. Introduction

Visual short-term memory (VSTM), or the capacity to temporarily retain a limited amount of visual information (Logie, 1989), is critical in our interactions with our environment and supports higher-order executive processes. Unfortunately, VSTM declines throughout the adult lifespan (Brockmole and Logie, 2013). Given the importance of VSTM in everyday functioning, a thorough understanding of the relation between VSTM and age is essential. A large number of studies have shown differential patterns of regional brain activation by memory load related to age, mainly expressed by greater activation in older adults, that can be interpreted by several accounts including the compensation-related utilization of neural circuits hypothesis (CRUNCH; Reuter-Lorenz and Cappell, 2008; but see Jamadar, 2020, for a conflicting finding and a systematic review). Regional activation studies typically report increased activity in the frontoparietal, dorsal attention, and salience networks and decreased activity in the default mode network in response to increased working memory load (e.g., Rottschy et al., 2012, Zuo et al., 2019). Modulation of activation by load is related to performance and is weakened with increasing age (Cappell et al., 2010; Heinzel et al., 2017; Kaup et al., 2014; Kennedy et al., 2017). While there is no doubt that frontoparietal regions are critical for VSTM, studies using multivoxel pattern analysis (MVPA) show that the contents of VSTM can be decoded from the visual cortex (e.g., Olmos-Solis et al., 2021) and that there is a decrease in classification performance as a function of load related to individual task performance (Emrich et al., 2013). It has been suggested that working memory maintenance arises from interactions between frontoparietal control regions and perceptual processing regions, where representations are maintained (e.g., Olmos-Solis et al., 2021; for an overview on this account see for example Eriksson et al., 2015; for a lifespan perspective see review Sander et al., 2012). Thus, functional connectivity between these regions is likely relevant for VSTM performance and examining age differences therein will inform our understanding of why working memory declines with age.

The effect of age on VSTM load-modulated functional connectivity is poorly understood. In young adults, VSTM load has been associated with within-network connectivity in the frontoparietal, dorsal attention, ventral attention, and visual networks (Eryilmaz et al., 2020; Liang et al., 2016; Soreq et al., 2019; Zuo et al, 2019), and between-network coupling among frontoparietal, dorsal attention, ventral attention and default mode networks (Eryilmaz et al., 2020; Liang et al., 2016). Furthermore, due to its role in sustaining attention, interactions between the dorsal attention network and other networks, is thought to be critical to memory performance at higher cognitive loads (Zuo et al., 2019).

A limited number of studies have investigated the effects of age on working memory load-modulated functional connectivity. Pongpipat and colleagues (2021) showed that functional connectivity within frontoparietal and default mode regions was strengthened as working memory load increased. With increasing load, the negative connectivity between the frontoparietal and default mode network also increased. Crucially, this pattern was invariant across the lifespan, but better performance only related to stronger negative coupling between these networks in middle-aged and older adults, not in younger adults or the oldest old. The youngest adults performed well, and the oldest adults poorly, regardless of connectivity between frontoparietal and default mode regions. The authors suggested that for adults in the middle and older age range, brain maintenance or compensation strategies are most important. In contrast, Heinzel et al. (2017) showed reduced working memory load-dependent modulation of connectivity between the dorsolateral prefrontal cortex and the inferior parietal lobe in older adults. Increasing this connectivity with increasing load was related to higher performance in older adults. A limitation of these studies is that they only included a very limited subset of brain regions. Crowell et al. (2020) used a whole-brain approach but only focused on one task network based on task-positive regions, including frontoparietal and sensorimotor regions, identified by a univariate analysis to be modulated by working memory load. They investigated network integration, defined as the ratio of connectivity within the task network versus between the task network and all other regions. While younger adults increased within-network connectivity with increasing load in the task network, older adults recruited a more distributed cortical network. This is in line with previous findings of reduced within-network specificity in older adults (Bethlehem et al., 2020; Chan et al., 2014; Geerligs et al., 2015) and the concept of age-related neural dedifferentiation (for a review, see Koen and Rugg, 2019).

Although these recent findings give some insight on how age affects load-modulated functional connectivity, a comprehensive investigation of the relationship between whole-brain connectivity in relation to VSTM load across the lifespan is missing. A whole-brain approach may be particularly critical, as a recent study showed that task-modulated connectivity not only involves task-active regions but also regions that are not activated or deactivated by the task (Di and Biswal, 2019). Thus, in the present study, we used a whole-brain approach to characterize age-related differences in load-modulated functional connectivity across the adult lifespan. Participants from the population-based Cambridge Centre for Ageing and Neuroscience (Cam-CAN) lifespan cohort (*N*=111, age 23-86) performed a visual short-term memory (VSTM) task during functional magnetic resonance imaging (fMRI). VSTM was measured with a delayed recall task for visual motion with three different loads. Whole-brain load-modulated connectivity was estimated using psychophysiological interactions in a hundred regions of interest, sorted into seven networks (Friston et al., 1997; Schaefer et al., 2018, Yeo et al., 2011). Using a delayed recall task, as opposed to an n-back task (which most previous studies have used), allowed us to compare task-related interactions during both encoding and maintenance periods. We expected: 1) increasing load to be associated with increased functional connectivity, particularly between attentional control and visual regions, 2) no effect of age on load-modulated functional connectivity between the frontoparietal and default mode network and decreased load-modulated connectivity between frontal and parietal regions with increasing age, and 3) better performance to be associated with higher load-modulated functional connectivity in networks modulated by VSTM load.

## 2. Materials and methods

### 2.1 Participants

A population-based sample of 280 healthy adults participated in Stage 3 of the Cam-CAN project. Participants completed a series of neuropsychological and neuroimaging experiments, of which the VSTM fMRI task is analyzed in this study. Half of the participants in Stage 3 were assigned to the VSTM task, of which 125 completed it. Eight participants were excluded based on poor performance on the lowest memory load trials of the task (single item, mean absolute error >30 degrees), two had incomplete EPI data, and four failed preprocessing of fMRI data. The final sample consisted of 111 participants with an age range of 23-86 years old (demographic information in Table 1, only grouped by age for illustrative purposes, not for analyses). MMSE scores ranged from 25 to 30. Eight participants had a missing MMSE score from the Stage 3 testing phase. Thus, for those participants, we used their MMSE score from Stage 1 for the means reported in Table 1 (the mean number of days between Stages 1 and 3 for these participants was 698 days, range 279-1317 days). Ethical approval for the Cam-CAN study was obtained from the Cambridgeshire 2 (now East of England – Cambridge Central) Research Ethics Committee. Participants gave written, informed consent before participating.

**Table 1.** Demographics

| Age group | Younger | Middle | Older |
| --- | --- | --- | --- |
| *N* | 36 | 35 | 40 |
| Age range (years) | 23-40 | 41-60 | 61-86 |
| Sex (male/female) | 18/18 | 16/19 | 23/17 |
| Handedness (right/left) | 33/3 | 35/0 | 38/2 |
| Highest education |  |  |  |
| University | 32 | 27 | 22 |
| A' levels | 1 | 5 | 11 |
| GCSE grade | 3 | 3 | 3 |
| None > 16 | 0 | 0 | 4 |
| MMSE *M* (*SD*) | 29.36 (1.02) | 28.91 (1.48) | 28.40 (1.34) |

Note. Sex based on self-report. Handedness = score on the Edinburgh Handedness Inventory (Oldfield, 1971). MMSE = Mini Mental State Examination.

### 2.2 Experimental design

The stimuli and task were adapted from Emrich et al. (2013) and have been described in detail in Mitchell et al. (in preparation). In short, VSTM was measured with a delayed recall task for visual motion with three different loads based on set size (Figure 1). Participants were asked to fixate on a dot in the center of a screen during a 7 second intertrial interval. Then three patches of coherently moving dots were presented sequentially, each in a different color. Patches were presented for 500 ms with a 250 ms blank interval. Depending on the set size, one, two or three patches displayed dots moving in a linear direction. Participants had to remember the motion direction and the corresponding color. The remaining patches showed dots moving in a circular motion at the same speed and could be ignored. During an 8 s blank interval, participants were to keep the motion direction(s) in mind. Finally, the probe display consisted of a circle with a line pointing to a random point on the circle, in one of the three colors that indicated which of the motion directions needed to be reported. Participants indicated the corresponding direction by rotating the line clockwise or counterclockwise by means of pressing keys on a button box. After confirmation of the direction by pressing a third key, or after 5 s, the next trial began.

**Figure 1.**
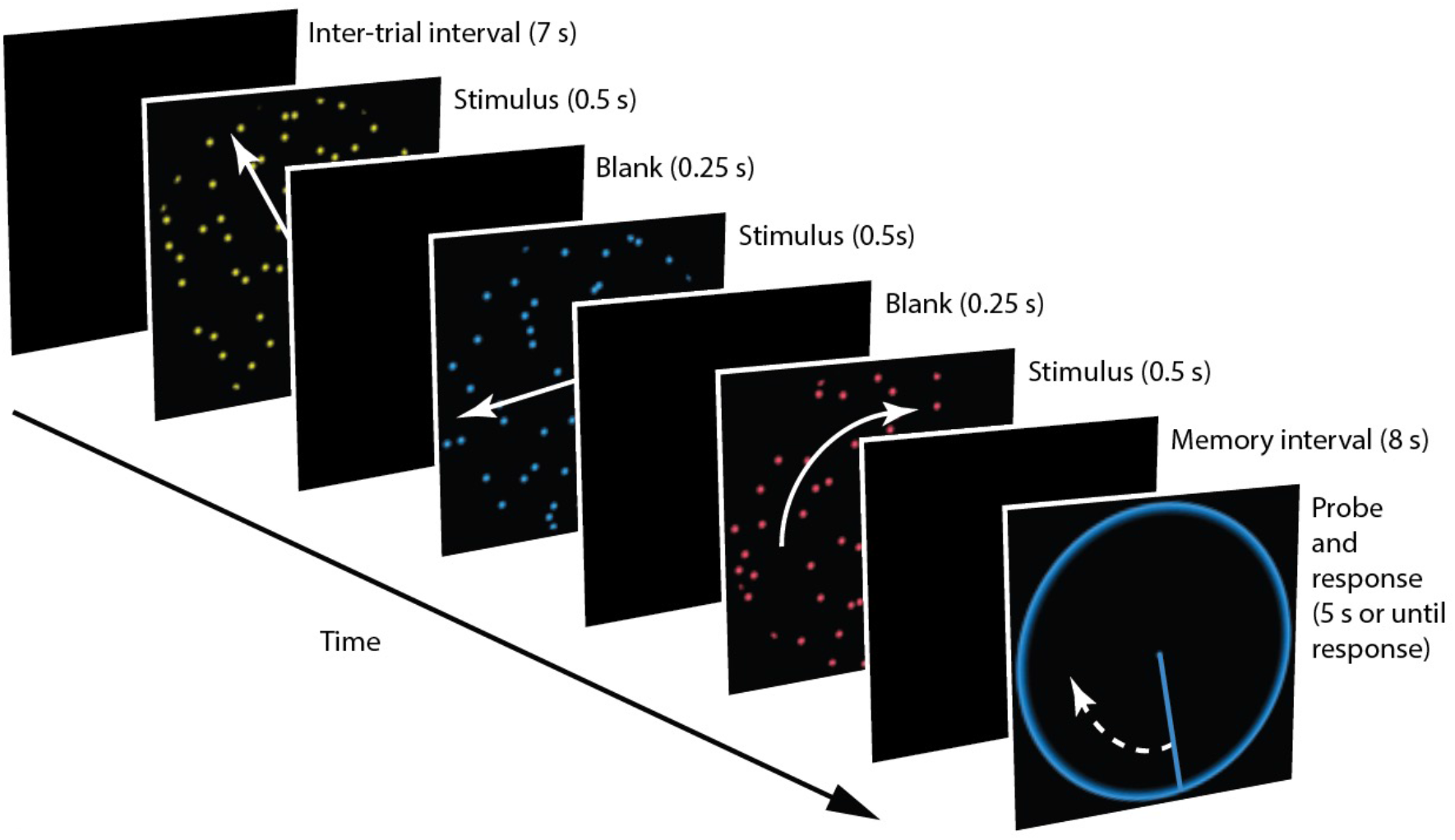
Example trial from the delayed-recall task. Participants were shown three patches of moving dots and were cued after a blank interval with a color of which the corresponding direction of motion needed to be reported.

The task consisted of 3 runs of 30 trials. Each run had 10 trials of each memory load (1, 2, and 3), counterbalanced with serial position (first, second, or third) and color (red, yellow, or blue). Trials within each run were presented in random order. On 27 trials per run, the motion direction of the probed patch was one of the three directions (7, 127, or 247 degrees). To prevent participants from noticing this pattern, random directions were used for the 3 remaining trials, as well as all the unprobed patches throughout the task (i.e., that needed to be memorized but were not tested).

### 2.3 Behavioral analysis

Response error, or the angular difference between the target value and the reported value, was used as a model-free measure of performance. The effects of load, age, and their interaction was analyzed using mixed effects modeling, with subject as random effect, and as fixed effects load and age. Analyses were conducted in R (version 4.2.0) using the lme4 package (Bates et al., 2015). Four models were built hierarchically such that effects were added one by one to see if additional predictors improved model fit (e.g., Sommet and Morselli, 2017). Predictors were added in the following order: subject, age, load, and age × load. Response error captures the mean absolute deviation from the target and was used instead of model estimates (such as precision; Zhang and Luck, 2008) because we found that model fit (log-likelihood) declined with age (Pearson correlation, set size 1 *r*(109) = -.36, *p* < .001, set size 2 *r*(109) < -.46, *p* < .001, set size 3 *r*(109) = -.53, *p* < .001).

Nevertheless, for completeness, we also report the results from a three-component mixture model (Zhang and Luck, 2008, modified by Bays et al., 2009) that distinguishes between different types of errors. This provides estimates of how many items (directions) were stored in memory (K), the precision of each item held in memory (kappa), the proportion of non-target errors (reporting a direction that corresponded to an unprobed patch), and the proportion of guess responses. The model described responses by a probabilistic mixture of errors centered on the target feature, centered on one of the non-target features, or distributed uniformly (guesses). Errors centered on the target or non-target features were described by a Von-Mises distribution (a circular analogue of the normal distribution). Further, because a previous study using the same dataset reported a significant response bias away from cardinal and towards oblique angles that increased with age Mitchell et al. (in preparation), we subtracted the mean response bias per direction from each response prior to running the mixture model. Finally, the correlation between different performance measures and age was analyzed per set size. Outliers on the response error measure were identified with a boxplot, based on a value higher than 1.5 times the interquartile range above the upper quartile (Tukey, 1977).

### 2.4 MRI acquisition

The MRI data were collected using a Siemens 3 T TIM TRIO system. During each of three functional runs while the participant was performing the task, T2*-weighted Echo Planar Images were acquired using the following parameters: Repetition time (TR) = 2 s, three echo times (TE = 12 ms, 25 ms and 38 ms), flip angle = 78 degrees, 34 axial slices of 2.9 mm thickness acquired in descending order with an inter-slice gap of 20%, in plane field of view (FOV) = 224 mm × 224 mm, and voxel-size = 3.5 mm × 3.5 mm × 3.48 mm. There were a variable number of scans per run, depending on reaction time, ranging from 294 to 349 (median 320).

A structural image was acquired for each participant, on a different day (median 49 days apart, range 75 to 1039). This used a T1-weighted 3D Magnetization Prepared Rapid Gradient Echo (MPRAGE) sequence with the following parameters: TR = 2250ms, TE = 2.98 ms, inversion time = 900 ms, 190 Hz per pixel, flip angle 9 degrees, FOV = 256 × 240 × 192 mm, and GRAPPA acceleration factor 2.

### 2.5 MRI Preprocessing

Imaging data were preprocessed with the automatic analysis (AA, release 3) batching system (https://imaging.mrc-cbu.cam.ac.uk/imaging/AA) using Statistical Parametric Mapping software (SPM12 release 7771; Wellcome Trust Centre for Neuroimaging, London, UK) and MATLAB (MATLAB version 9.10.0.1739362 (R2021a). Natick, Massachusetts: The MathWorks Inc.). The structural images were rigid-body registered with an MNI template brain, bias corrected, segmented, and warped to match a study-specific anatomical template based on the whole CamCAN Stage 3 sample using the DARTEL procedure (Ashburner, 2007; Taylor et al., 2017), which was subsequently normalized into MNI space. The functional images were then realigned, corrected for slice-time acquisition, rigid-body coregistered to the structural image, transformed to MNI space using the warps and affine transforms from the structural image, smoothed (8 mm FWHM Gaussian kernel), and resliced to 3 × 3 × 3 mm voxels.

The first-level GLM for each participant comprised three neural components per trial: (1) encoding, modeled as an epoch of 2 s duration at onset of the first moving dot pattern; (2) maintenance, modeled as an epoch of 8 s from the onset of the blank screen; and (3) probe, modeled as an epoch of the duration from the onset of the probe till response. Each of these components were modeled separately for the three memory load levels. Two block regressors were added to model the three different runs.

### 2.6 Correlational PPI

Whole-brain functional connectivity matrices were estimated for the contrast set size 3 > set size 1, for each subject by using a correlational psychophysiological interaction (cPPI) approach as implemented by the cPPI toolbox (Fornito et al., 2012; Friston et al., 1997). Since we had no a priori predictions about directionality we used cPPI which is based on partial correlations between ROIs to isolate task-related changes in functional coupling between regions, resulting in symmetrical connectivity matrices. This avoids the arbitrary distinction between seed and target as is required in traditional PPI (Wang et al., 2018). We obtained connectivity terms for 100 ROIs corresponding to the cortical Schaefer parcellation (Schaefer et al., 2018). The ROIs were sorted into seven networks from Yeo et al. (2011): visual, somatomotor, dorsal attention, ventral attention, limbic, frontoparietal, and default mode. As the “limbic network” only consisted of the orbitofrontal cortex and temporal pole, we used the term OFC-TP to refer to this network for the sake of clarity. Within- and between-network connectivity was defined based on these networks. The PPI terms were obtained by modeling the task effects, the physiological signal, and their interaction. For each participant, time series were extracted from each ROI based on the first-level GLM design matrix. For each pair of ROIs, the time courses of both ROIs were deconvolved as implemented in SPM (Gitelman et al., 2003). Both deconvolved time courses are multiplied by the unconvolved task regressors for the contrast of interest (set size 3 > set size 1) to generate the deconvolved PPI terms. These PPI terms were then reconvolved. Finally, the partial correlation between these two PPI terms was computed adjusting for: the raw time courses of the two ROIs, the task regressors creating the contrast of interest, the time-courses of the remaining 98 ROIs, and nuisance signals. Thirty-four nuisance regressors were added, representing six motion parameters estimated in the realignment stage, time-series from white matter tissue and cerebrospinal fluid (CSF), their derivatives, plus quadratic functions of those 16 parameters. Masks for white matter and CSF were based on the FieldMap toolbox in SMP12 (probability maps were thresholded at .8 and .5 respectively, based on visual inspection for no overlap with the Schaefer cortical atlas). This was done for all ROI pairs, resulting in a 100 × 100 symmetrical matrix for encoding and maintenance separately.

### 2.7 Group-level analysis

We first ran, for the partial connectivity between each pair of ROIs, a one-sample t-test for the effect of task load for the contrast set size 3 > set size 1, with a threshold of r > .5 to identify which ROI pairs showed a medium to large effect of memory load.

Subsequently, per ROI pair, we ran three GLMs, in a stepwise manner, to assess the relationship between age, performance, and the age by performance interaction and brain-wide load-modulated connectivity. For performance we used response error from set size 3, as using the difference score between set sizes 3 and 1 would be noisier and performance at set size 3 is most sensitive to individual differences. Performance and age were mean-centered before calculating the interaction term. In the first model we tested the effect of age on the load-dependent changes in connectivity (cPPI ∼ age). In the second model we added performance as a regressor to test the effect of performance on modulation of connectivity by load, while controlling for age (cPPI ∼ age + performance). The third model tested for the interaction between age and performance (cPPI ∼ age * performance). As effect sizes of associations between neural measures and cognition are small (Marek et al., 2022) and our whole-brain approach necessitated correction for many tests, we repeated the analyses of the second and third model restricted to network pairs where load-modulated functional connectivity was significant in at least 50% of the edges (ROI-pairs) in either the encoding or maintenance connectivity matrix.

Both the one-sample t-test and the 3 GLM models were controlled for the family-wise error rate (FWE) using the Network Based Statistics (NBS) toolbox (Zalesky et al. 2010), with a threshold of 2.361 (*t*-value corresponding to a *p*-value of .01 with a DF of 110) and *p* < 0.025 (2 comparisons, positive and negative contrast) graph component significance (equivalent to a cluster in mass univariate analysis, in this context a data-driven network of related edges) as determined by permutation testing (n = 5000) using the method of Freedman and Lane (1983). Results are reported as the number of significant edges. NBS has been found to offer more power than FDR correction for identifying distributed networks of edges (Fornito et al., 2013) and has been used to identify functional networks associated with memory performance and age before (Capogna et al., 2022). BrainNet Viewer was used for visualisation of networks on a 3D-brain (Xia et al. 2013, http://www.nitrc.org/projects/bnv/).

### 2.8 Data availability

Derived neuroimaging data, behavioral data, and code are available in the Open Science Framework at https://osf.io/w3s74/. Raw data can be requested from the Cam-CAN access website (https://camcan-archive.mrc-cbu.cam.ac.uk/dataaccess/). For a complete description of Cam-CAN data and pipelines, see Shafto et al. (2014) and Taylor et al. (2017).

## 3. Results

### 3.1 Behavioral results

Using mixed linear models, we analyzed the effects of age and VSTM load on response error. The first model contained only subject as random effect and confirmed that accuracy across memory loads was correlated within individuals (ICC = .40). Adding age to the model improved the model fit (χ2(1)=28.53, p < .001), as did load (χ2(1)=173.14, p < .001), and the age × load interaction (χ2(1)=41.58, p < .001). Pairwise contrasts showed that response error on set size 1 was different from that of set size 2 and set size 3, and there was a difference between set size 2 and set size 3 (*p* < .001, Figure 2A). The positive correlation between age and response error is significant at all loads (Table 2), but as can be seen in Figure 2B, the effect of age was more pronounced at higher loads. As the boxplot identified one outlier on response error at set size 3 (i.e., our outcome measure of interest for fMRI analyses), we reran all analyses excluding this subject (age 54, female). This did not lead to different results.

**Table 2.**
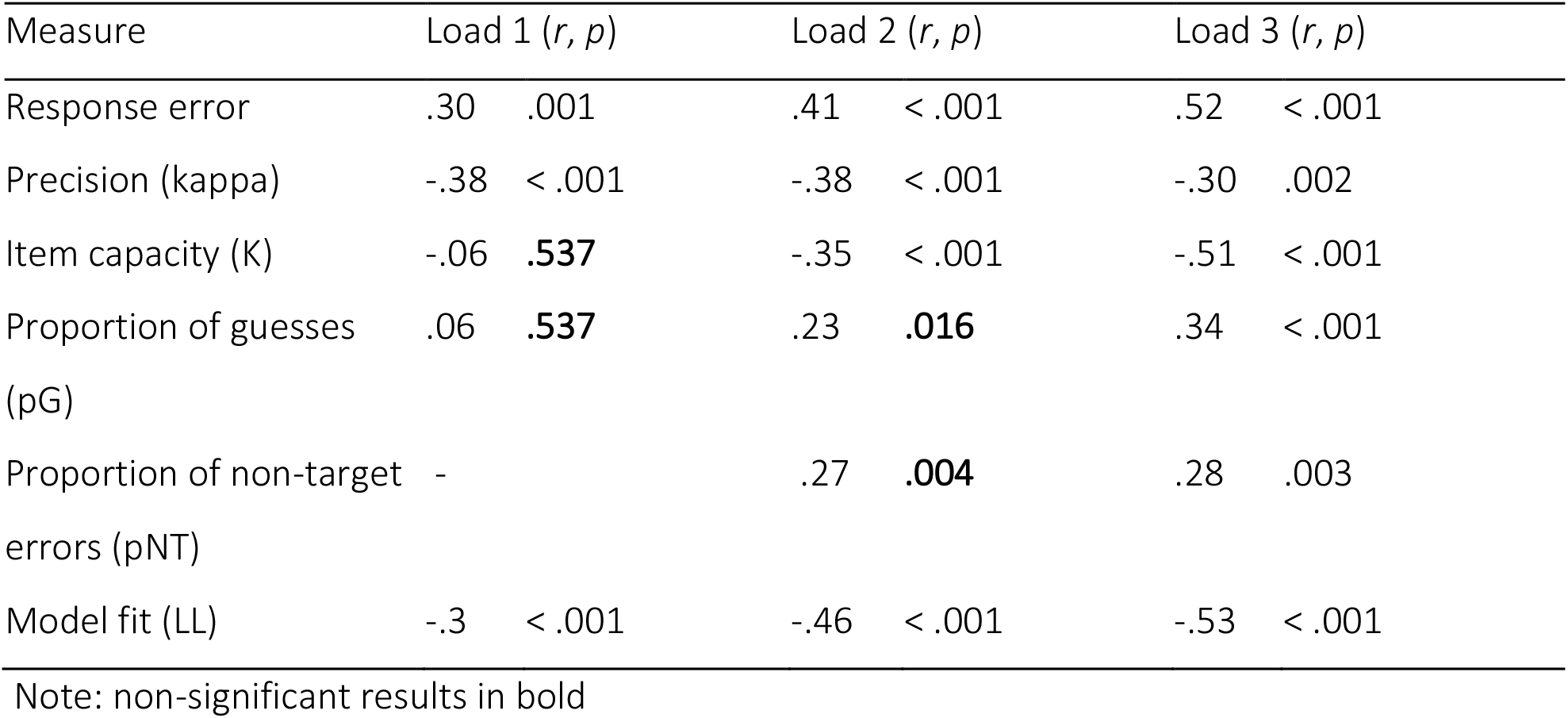
Correlations between VSTM performance measures and age per memory load

**Figure 2.**
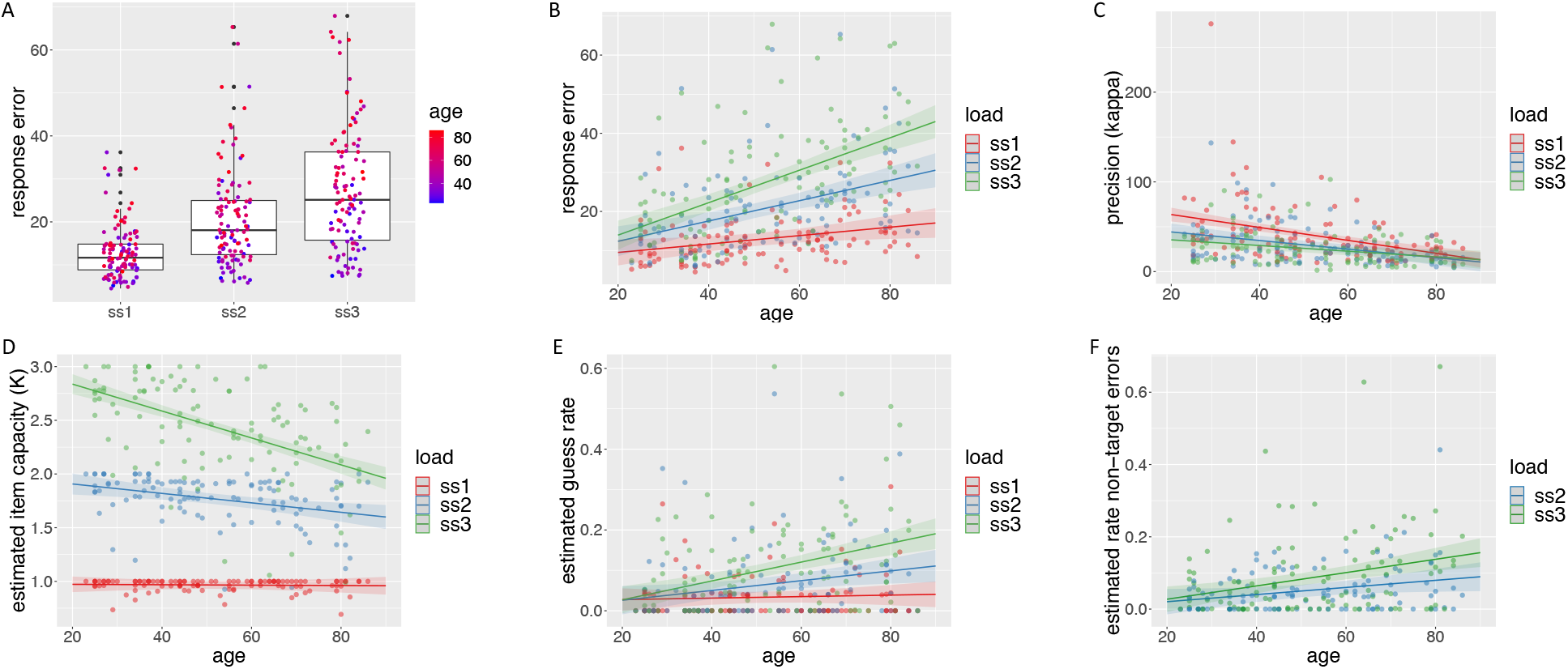
Effects of memory load and age on VSTM performance. A) response error shown as boxplot per load with age on a color scale from young in blue to old in red. In panels B to F, the linear regression line for the effect of age is shown per set size, along with its 95% confidence interval, for the B) response error, C) precision (kappa), D) item capacity (K), E) estimated guess rate, and F) estimated rate of non-target errors.

To evaluate the contribution of different sources of error, we applied a three-component mixture model (Bays et al., 2009). Most of these measures correlated significantly with age (Table 2, Figure 2C-F), except for estimated item capacity at set size 1, guess rate at set size 1 and 2, and proportion of non-target errors at set size 2 (adjusted alpha .05 / 14 = .0036; note that non-target errors are not possible for set size 1). A decrease in precision, and an increase in guess rate and non-target errors, all contribute to a higher response error with increasing age at a high memory load. However, as already mentioned, increasing age was associated with a lower model fit at all three loads and thus, we decided to use raw response error in relating performance to functional connectivity.

### 3.2 Whole-brain functional connectivity

Calculating whole-brain load-modulated functional connectivity for the contrast of load (set size 3 > set size 1), we assessed the effect of memory load, and how this modulation by load is associated with age, performance, and their interaction.

#### 3.2.1 Main effect of load

Load-modulated functional connectivity strengthened with increased positive connections at higher load both within and between networks (Figure 3 A and D). An NBS-based one-sample t-test with a threshold of r > .5 was applied to identify edges that showed a medium to large effect of load (FWE-corrected, number of significant edges for encoding = 474, and maintenance = 646, Figure 3 B and E). The strongest within-network connectivity was in the dorsal attention network followed by the visual network, and to a lesser extent the somatomotor, ventral attention, and frontoparietal networks (see Figure 3 C and F, which show the proportion of edges that survive thresholding per network). Between-network connectivity increased primarily between the somatomotor, dorsal attention, and ventral attention networks, and between the dorsal attention and the frontoparietal networks. The pattern of increased positive correlations with increasing load was highly comparable for encoding and maintenance^1^.

**Figure 3.**
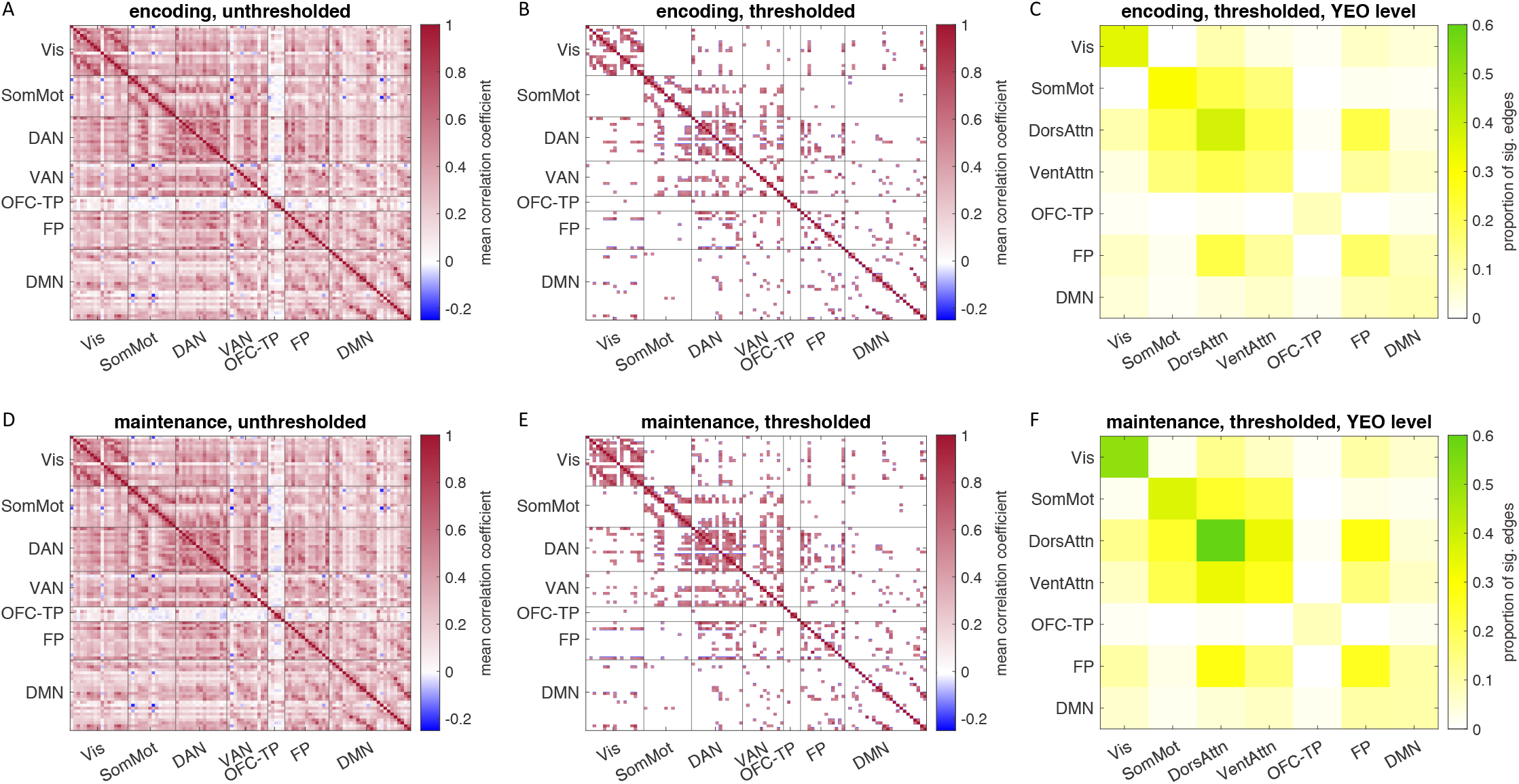
Partial correlation coefficient matrices for mean encoding (A and B) and maintenance (D and E) connectivity for the contrast set size 3 > set size 1. ROIs are sorted into seven networks from Yeo et al. (2011): visual (Vis), somatomotor (SomMot), dorsal attention (DAN), ventral attention (VAN), orbitofrontal-temporal pole (OFC-TP), frontoparietal (FP), and default mode (DMN). Red colors indicate a positive partial correlation, while blue colors indicate a negative partial correlation. Panels A and D show the unthresholded matrices. Panels B and E show the results of a one-sample t-test for a medium to large effect size (thresholded r > .5, FWE-corrected *p* < .025). Panels C and F show the proportion of edges that survive thresholding per network cell, with green indicating a higher proportion.

#### 3.2.2 Effect of age

With increasing age, there was a significant decrease in load-modulated connectivity differences between set size 1 and set size 3 throughout the brain. Figure 4 A and D show the correlation between age and load-modulated connectivity for each ROI pair (number of significant edges based on NBS for encoding = 2449 and maintenance = 2673, Figure 4 B and E). Age-related declines were most pronounced within the somatomotor, frontoparietal, and default mode networks, and between the visual network and the dorsal attention and frontoparietal networks (Figures 4C and 4F show the proportion of significant edges per network pair). During maintenance there was additionally a pronounced decline in load-modulated functional connectivity within the dorsal and ventral attention network, and between the visual network and the somatomotor and ventral attention networks, and the dorsal attention and frontoparietal networks. There were no significant positive effects of age (graph component FWE-corrected *p* = .11 and *p* = .17 for encoding and maintenance respectively); that is, no connections showed a load-dependent increase in connectivity that was stronger in older compared to younger adults. Older adults do still show positive load-modulated functional connectivity but to a lesser extent compared to younger adults, as illustrated in Figure 5 for average within network connectivity by age group (young: 20-40; middle: 41-60; old: older than 60 years).

**Figure 4.**
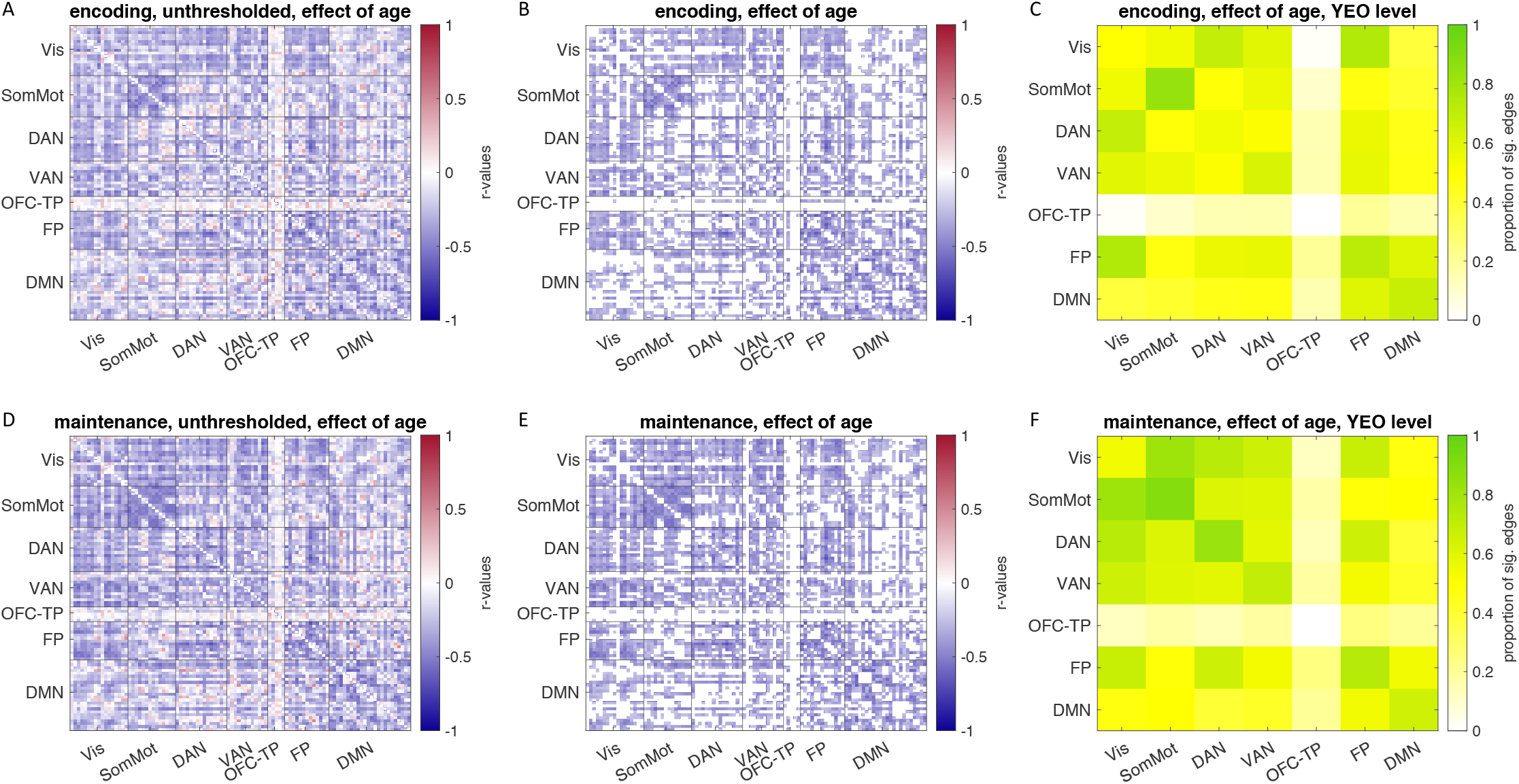
Correlation matrices for encoding (A and B) and maintenance (D and E) for the correlation between age and load-modulated connectivity strength. Red colors indicate a positive correlation, while blue colors indicate a negative correlation. Panels A and D show the unthresholded matrices. Panels B and E show only edges where the effect of age is significant based on an NBS based t-test (FWE-corrected, *p* < .025). Panels C and F show the proportion of edges that show a significant effect of age per network cell, with green indicating a higher proportion.

**Figure 5.**
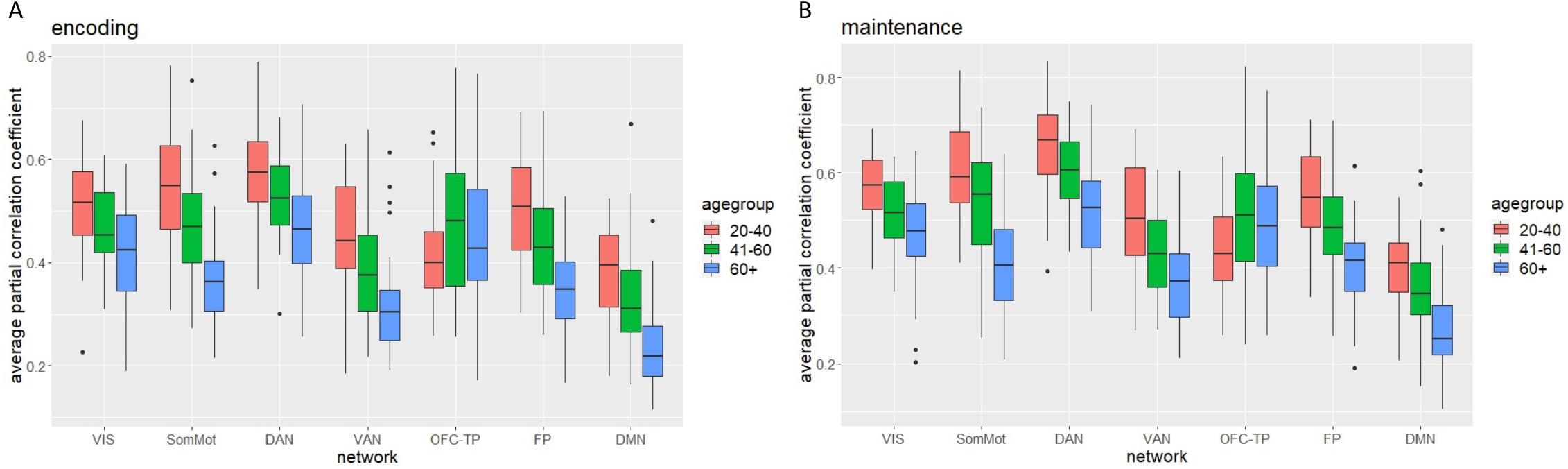
Average load-modulated connectivity strength (partial correlation coefficient) within each network per age group. Participants are split into young (20-40 years old), middle (41-60 years old), and older (60+) adults for illustrative purposes. Left during encoding, right during maintenance.

#### 3.2.3 Effect of Performance

Running the model (predicting connectivity from performance, controlling for age) on the full matrices for encoding and maintenance did not result in any effects of performance that survived correction. The two networks that were most modulated by effect of task load were the visual and dorsal attention networks (Figure 3 C and F). Running the model on just these two networks showed positive and negative associations between performance and connectivity within the visual network (which did not survive correction), and negative associations in the dorsal attention network (Figure 6 A and D). After thresholding, lower response error (i.e., better performance) was associated with a significant increase in load-modulated connectivity strength within the dorsal attention network, controlling for the effect of age (number of significant edges for encoding = 17, and maintenance = 22, Figure 6 B and E; scatterplot of the relationship between average load-modulated connectivity in the dorsal attention network and performance controlled for age in supplementary materials S2, Figure S2.1). This included bilaterally the posterior and superior parietal lobe, the precentral ventral frontal cortex, and the frontal eye fields. Additionally, in the left hemisphere, connectivity in the lateral parietal cortex (Figure 6 C and F).

**Figure 6.**
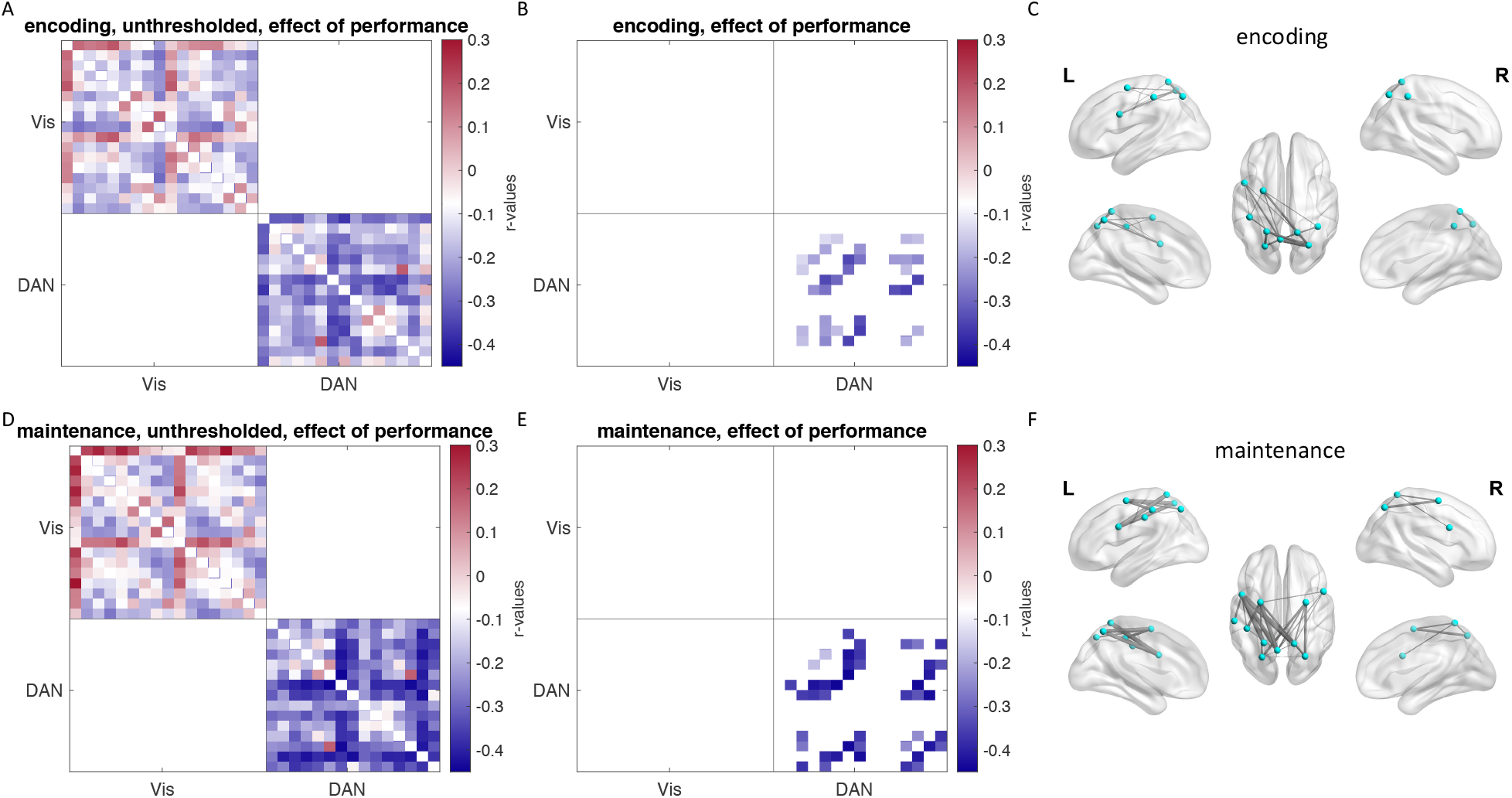
Correlation matrices for encoding (A and B) and maintenance (D and E) for the correlation between performance (response error at set size 3) and load-modulated connectivity strength. Red colors indicate a positive correlation, while blue colors indicate a negative correlation. Panels A and D show the unthresholded matrices. Panels B and E show only edges where the effect of performance is significant based on an NBS based t-test (FWE-corrected, *p* < .025). Panels C and F show the significant edges overlaid to a 3D-brain.

While the effect of memory load was most pronounced within the visual and dorsal attention networks, other within and between network connections also survived thresholding for the effect of load (see Figures 3 B and E). In a further, exploratory analysis, we therefore tested whether connectivity is associated with performance when including all surviving edges for the effect of load. This analysis replicated our previous results, showing that most edges related to performance were within the dorsal attention network, with an additional a few in the frontoparietal network, and between these networks (supplementary materials S2, Figure S2.2). However, the results did not survive correction for testing positive and negative effects separately (*p*<.025; encoding *p* = .031 maintenance *p* = .033). The number of edges is different in both analyses (241 within the visual and dorsal attention network versus 474 and 646 edges for the effect of load during encoding and maintenance, respectively).

#### 3.2.4 Interaction effect of Age × Performance

Last, we tested the interaction effect of age × performance over all edges and restricted to the visual and dorsal attention networks. This yielded no significant results.

## 4. Discussion

In this study, we investigated 1) VSTM load-modulated functional connectivity across the cortex during encoding and maintenance, 2) age differences therein, and 3) how load-modulated functional connectivity relates to performance. Connectivity was stronger for high compared to low VSTM load, especially within the dorsal attention and visual networks, and between the somatomotor, dorsal attention, and ventral attention networks, as well as between the dorsal attention and frontoparietal networks. Age was associated with a reduction in load-modulated connectivity across the cortex. The effect of age was most pronounced within the frontoparietal, default mode, and somatomotor networks, and during maintenance additionally within the dorsal and ventral attention networks and between the visual network and other networks. Finally, increased load-modulated connectivity within the dorsal attention network was related to better performance on the task, in an age-invariant way. Together, these results contribute to a better understanding of working memory decline with age.

### 4.1 VSTM load-modulated functional connectivity

Connectivity strength was higher for high versus low working memory load. We observed the strongest load-modulated connectivity within the dorsal attention and visual networks, both during encoding and maintenance. It is important to note that increased connectivity within the visual network with increasing load cannot be easily explained by more visual input, as the amount of visual input and average motion per item was matched at all three set sizes. Load-modulated connectivity was highly comparable between the encoding and maintenance phases. This was also demonstrated in a previous study using a visuospatial working memory task (Soreq et al., 2019) and is in line with the sensory recruitment theory. According to the sensory recruitment account of working memory (e.g., D’Esposito and Postle, 2015), the same cortical regions involved in perceptual processing are recruited for maintenance during working memory. Compelling evidence for this account derives from studies using MVPA that show that the visual cortex supports representations of visual features during the delay period, despite mean activity in the visual cortex dropping to baseline levels during that same period (reviewed in D’Esposito and Postle, 2015; and in Teng and Postle, 2021). A study by Emrich et al. (2013), showed that with increasing memory load, MVPA decoding performance in the visual cortex declined, along with delayed-recall precision. In contrast, while the intraparietal sulcus shows load-dependent delay-period activity, working memory content could not be readily decoded from this region at higher loads (Emrich et al., 2013). The parietal cortex is thought to be involved in attentional control needed for context-binding, prioritization of items, and resistance to distraction (Gosseries et al., 2018; Teng and Postle, 2021). The function of the frontoparietal network, according to the sensory recruitment framework, is top-down control and abstract representations (Scimeca et al., 2018). Our results show a moderate modulation by load in functional connectivity within the frontoparietal network, which may reflect the need for more top-down control at a higher load.

We also found strong load-modulated connectivity within the somatomotor network, which was also observed by Soreq and colleagues (2019) using a visuospatial change detection task. This might be related to the spatial aspect of the task used here. Participants likely maintained a spatial representation of each motion direction in order to respond correctly. Connectivity within the somatomotor network has been associated with prospective motor coding during a spatial delayed recall task (Purg et al, 2022), with greater connectivity seen when there is a predictable relation between the motor response and the location of the memorized target (motor response from center to target > motor response from random location to target). Prospective motor coding can be used to execute a spatial response but also to support rehearsal and evaluate a memory probe (Postle, 2006). To stress the coupling between sensory maintenance and motor intention, D’Esposito and Postle (2015) argue for the use of the label “sensorimotor recruitment” instead of the more common label “sensory recruitment” models. In sum, our finding of moderate to strong load-dependent changes in functional connectivity within the dorsal attention, visual, frontoparietal, and somatomotor networks is in line with the sensory recruitment account of VSTM maintenance.

We also observed load-modulated changes in connectivity *between* networks, including the dorsal attention, ventral attention, and somatomotor networks, and between the dorsal attention and frontoparietal networks. The functions of these networks include visuospatial attention, processing salience, somatosensory and motor processing, and working memory and inhibition (respectively; reviewed in Uddin et al., 2019). Higher working memory load plausibly results in stronger connectivity between these networks as more top-down control is required. Interestingly, load-modulated connectivity between the visual and dorsal attention networks was limited. It might be that functional connectivity between those networks is consistent across different memory loads. A second possible explanation is that between network connectivity is only modulated by load during the response phase when a comparison is made between the memorized items to select a response (Xu, 2017).

Our finding of stronger connections within the visual and dorsal attention networks at higher load aligns with previous studies that used a visual working memory task (Zuo et al., 2019; Soreq et al., 2019). A second finding consistent with previous work is increased connectivity within the frontoparietal network at higher working memory load (Eryilmaz et al., 2020; Liang et al., 2016; Pongpipat et al., 2021; Soreq et al., 2019; but see Zuo et al., 2019). Two other studies investigated connectivity within the somatomotor network: using a Sternberg task with letters, Eryilmaz et al. (2020) did not find connectivity within the somatomotor network contributing to classification of different memory loads, while Soreq et al. (2019), who used a visuospatial task, did find positive load-modulated connectivity within the somatomotor network. This fits with our interpretation that load-modulated connectivity within the somatomotor network might be related to the spatial aspect of the task. Increased connectivity between the dorsal and ventral attention network and the frontoparietal network with increased load is consistent across different types of tasks (Eryilmaz et al., 2020). The main discrepancy between our results and previous studies is the limited effect of working memory load on connections within the default mode network and connectivity between this network and other networks (Eryilmaz et al, 2020; Liang et al., 2016; Pongpipat et al., 2021). Pongpipat et al. (2021) reported an increase in negative connectivity between the frontoparietal and default mode network as load increased. This effect was stronger when comparing task with a control condition than for the linear effect of load. Further, a study by Zuo et al. (2019), showed that while the default mode network was strongly associated with performing an N-back, this was to a similar extent in the contrasts of baseline vs 0-back, and 0-back vs 2-back. It might be that we did not find negative load-modulated connectivity of the default network because we did not compare to a no-load baseline.

### 4.2 Age effects on load-modulated functional connectivity

We found negative effects of age on the modulation of functional connectivity by load in widespread networks throughout the cortex. Older adults showed less increase in connectivity as task demands increased. Reduced connectivity with age in the default mode network is a consistent finding in the literature across both task-based and resting state studies (Andrews-Hanna et al., 2007; Bethlehem et al., 2020; Damoiseaux et al., 2008; Geerligs et al., 2012, 2015; Grady et al., 2010; Sambataro et al., 2010; Samu et al., 2017; Tsvetanov et al., 2016), one that we replicate here. Our study additionally shows a novel finding of decreased load-modulated connectivity with increasing age within the somatomotor, dorsal attention, ventral attention, and frontoparietal network, and between the visual network and other networks. Decreases in within and between network connectivity across the cortex with increasing age have been reported in a language task (Zhang et al., 2021) and within networks supporting higher level cognitive functions during resting-state (Geerligs et al., 2015).

The resting-state literature describes decreased segregation of functional networks with increasing age, with weaker within network connectivity and greater connectivity between networks (Chan et al., 2014; Geerligs et al., 2012, 2015). Our result of weaker load-modulated connectivity with increasing age within and between all networks except the orbitofrontal-temporal pole network, show that these findings from resting-state data do not easily translate to task-modulated connectivity. This stresses the relevance of a task-based approach for understanding how age affects cognitive functioning (Campbell and Schacter, 2016).

A possible explanation for reduced modulation in network functional connectivity by working memory load in older adults, is that they may already be close to ceiling in terms of resources at set size one. We found that older adults do still show positive load-modulated connectivity (Figure 5), but to a lesser extent than younger adults. Possibly our lowest load condition (which required remembering a single direction while ignoring rotating dots in two other colors) was more demanding than that of some other studies using, for example, a 0-back task. In a previous study, older adults showed a large change in functional connectivity moving from baseline to a simple task, while connectivity was comparable when moving from a simple task to a more demanding task; in contrast, younger adults showed the largest change from a simple to a more demanding task (Geerligs et al., 2013). This explanation fits well with our finding that, across the cortex, younger adults showed a larger increase in functional connectivity from set size one to set size three. If older adults were already expending close to maximum resources at set size one, they would not show much further increase with task load (compatible with CRUNCH; Reuter-Lorenz and Cappell, 2008).

For the maintenance phase specifically, we found a strong negative effect of age on within network connectivity in the dorsal attention network. This network has been associated with (maintenance of) visuospatial attention and top-down selection for stimuli and responses (Uddin et al., 2019). Older adults especially show a decline in attentional control (Bier et al, 2017; Hasher and Zacks, 1988; Kato et al., 2016), and a lessened ability to inhibit concurrent distraction (e.g., May, 1999; Campbell et al., 2012) and previously attended (but no longer relevant) information (e.g., Henderson et al., 2020; Healey et al., 2013; Sylvain-Roy et al., 2015; for a recent review, see Campbell et al., 2020). Older age in our study was related to greater response error, which at the highest load, the mixture model suggests was due to lower precision, lower item capacity, and a higher rate of non-target errors and guess responses. Lower precision and item capacity suggests an age-related decline in the ability to maintain multiple high-resolution representations, while a higher rate of non-target errors and guesses might relate to a decline in inhibition and attentional control. A pronounced negative effect of age on load-modulated connectivity between the visual network and other networks might also be related to different use of strategy, with older adults relying less on fine-grained visual representations and more on verbal labels (Forsberg et al., 2019).

### 4.3 Performance effects on load-modulated functional connectivity

We found that individuals with a lower response error (i.e., better performance) are those who show a higher increase in functional connectivity with increasing load within the dorsal attention network, while controlling for the effect of age. The dorsal attention network is primarily associated with visuospatial attention. Various models of memory have indicated attention as crucial for working memory (Baddeley and Hitch, 1974; Cowan, 1988). As working memory representations are vulnerable to interference, sustained attention is crucial both to maintain relevant information and to ignore non-relevant stimuli (e.g., Vogel et al., 2005). With increasing load, more attentional control is needed as more items need to be maintained in memory, one of which must be selected at test while inhibiting the others. Our finding that the visual and dorsal attentional networks are most sensitive to load but only modulation in connectivity in the dorsal attention network is significantly related to performance, converges with a large body of research which suggests that individual differences in working memory performance are determined for a large part by variability in attentional control (for a review, see Erikson et al., 2015).

Our statistical testing for an interaction effect revealed no evidence that the relationship between performance and load-modulated connectivity differed with age. This suggests that increased connectivity in the dorsal attention network with increasing load relates to better performance in a similar way across the lifespan.

### 4.4 Limitations and methodological remarks

In this study we investigated the modulation of functional connectivity by working memory load. There are many confounds when studying age-related differences in functional connectivity, like head motion (Geerligs et al., 2017), vascular health (Tsvetanov et al., 2020), and motor skill for task-based connectivity. By using a contrast for two different task loads, we obtained a measure that showed the difference within each individual between two conditions. This within-individual measure is less sensitive to between-individual confounds (Geerligs et al., 2017), and demonstrates that the difference in connectivity from low to high load is smaller in older adults. However, this does not give information on whether older adults have higher connectivity strength at low load compared to younger adults, as CRUNCH would predict (discussed above; Reuter-Lorenz and Cappell, 2008), and/or lower connectivity strength at high load. We could not test this here due to our lack of a zero-load condition in the task. Thus, it is important to keep in mind when interpreting these findings that an ROI or network pair that does not show a modulation by task load might still change in strength of connectivity between rest and task.

A second limitation is that the patterns of connectivity during encoding and maintenance cannot fully be dissociated. The encoding period always preceded maintenance, and as is typical for most working memory paradigms, the delay-period had a fixed length without jitter (Crowell et al., 2020). We included regressors for the events that we were not interested in for each specific contrast (e.g., regressors for the encoding and response periods when examining the maintenance phase), but this had no influence on the connectivity matrices (i.e., partial correlations did not change). Thus, in the current design we cannot exclude the possibility of some contamination from the encoding period during maintenance. A related issue typical for working memory task designs, is the difference in duration of the encoding and maintenance period. A longer period might have a better signal-to-noise ratio and therefore, result in stronger effects (which is indeed what we found - see supplementary materials). However, with a closer look at the effects of age and performance on connectivity patterns, there are slightly different patterns of connectivity during encoding and maintenance. Specifically, stronger effects of age between the frontoparietal and the visual and default mode network were observed during encoding. This suggests that the design is to some extent able to detect differences between the two task periods, and that the highly similar patterns are not just due to spill-over effects.

Third, the advantage of a whole-brain approach is that we were able to demonstrate how widespread the effect of age is on load-modulated functional connectivity. However, it does come at the cost of multiple comparisons. Whereas effects of age were clearly detectable, the effects of cognition on neural signals are often weaker (see for a review on brain-wide association studies Marek et al., 2022), made worse here by the need to control for age.

Therefore, we focused on the effect of performance only in the network cells that showed the strongest effect of task load. We were able to demonstrate that an increase in connectivity strength within the dorsal attention network was related to better performance, while controlling for age. It might be that functional connectivity within and between other regions contributed to performance too. Our exploratory analysis on all above-threshold connections suggested that in addition to within network load-modulated connectivity in the dorsal attention network, connections within the frontoparietal network and between the frontoparietal and dorsal attention network might be relevant to performance. This should be investigated in future studies using an even larger sample.

## 5. Conclusion

The current study confirms that functional connectivity strengthens in response to increasing working memory load. This was especially true in the dorsal attention and visual networks during encoding and maintenance. A novel finding was that VSTM load-modulated functional connectivity decreased with age across most of the cortex both within and between networks, possibly because older adults are already close to ceiling in terms of resources at the lowest load. In the dorsal attention network, increased load-modulated functional connectivity was related to better VSTM performance. Notably, the patterns of connectivity related to performance were age-invariant. These findings are in line with the sensory recruitment model of working memory and stress the role of attentional control in visual working memory across the adult lifespan.

## CRediT roles

**Selma Lugtmeijer:** Conceptualization, Data curation, Formal analysis, Methodology, Project administration, Visualization, Writing - original draft, **Linda Geerligs:** Conceptualization, Supervision, Writing - review and editing, **Kamen A. Tsvetanov:** Methodology, Writing - review and editing, **Daniel J. Mitchell:** Conceptualization, Data curation, Formal analysis, Methodology, Software, Writing - review and editing, **Cam-CAN:** Investigation, Data curation, **Karen L. Campbell:** Conceptualization, Funding acquisition, Project administration, Resources, Supervision, Writing - review and editing.

## Acknowledgements

The Cambridge Centre for Ageing and Neuroscience (Cam-CAN) was supported by the UK Biotechnology and Biological Sciences Research Council (grant number BB/H008217/1), together with support from the UK Medical Research Council Cognition & Brain Sciences Unit (CBU) and University of Cambridge, UK. The authors are grateful to the CamCAN respondents and their primary care teams in Cambridge for their participation in this study. The authors also thank colleagues at the MRC Cognition and Brain Sciences Unit MEG and MRI facilities for their assistance. The Cam-CAN corporate author consists of the project principal personnel: Lorraine K Tyler, Carol Brayne, Edward T Bullmore, Andrew C Calder, Rhodri Cusack, Tim Dalgleish, John Duncan, Richard N Henson, Fiona E Matthews, William D Marslen-Wilson, James B Rowe, Meredith A Shafto; Research Associates: Karen Campbell, Teresa Cheung, Simon Davis, Linda Geerligs, Rogier Kievit, Anna McCarrey, Abdur Mustafa, Darren Price, David Samu, Jason R Taylor, Matthias Treder, Kamen A Tsvetanov, Janna van Belle, Nitin Williams, Daniel Mitchell, Simon Fisher, Else Eising, Ethan Knights; Research Assistants: Lauren Bates, Tina Emery, Sharon Erzinçlioglu, Andrew Gadie, Sofia Gerbase, Stanimira Georgieva, Claire Hanley, Beth Parkin, David Troy; Affiliated Personnel: Tibor Auer, Marta Correia, Lu Gao, Emma Green, Rafael Henriques; Research Interviewers: Jodie Allen, Gillian Amery, Liana Amunts, Anne Barcroft, Amanda Castle, Cheryl Dias, Jonathan Dowrick, Melissa Fair, Hayley Fisher, Anna Goulding, Adarsh Grewal, Geoff Hale, Andrew Hilton, Frances Johnson, Patricia Johnston, Thea Kavanagh-Williamson, Magdalena Kwasniewska, Alison McMinn, Kim Norman, Jessica Penrose, Fiona Roby, Diane Rowland, John Sargeant, Maggie Squire, Beth Stevens, Aldabra Stoddart, Cheryl Stone, Tracy Thompson, Ozlem Yazlik; and administrative staff: Dan Barnes, Marie Dixon, Jaya Hillman, Joanne Mitchell, Laura Villis.

Further, Linda Geerligs was supported by a VIDI grant of the Netherlands Organization for Scientific Research (grant number VI.Vidi.201.150). Daniel J. Mitchell was supported by the Medical Research Council intramural program SUAG/085.G116768. Kamen A. Tsvetanov was supported by the Guarantors of Brain (G101149); and Karen L. Campbell by the Natural Sciences and Engineering Research Council of Canada (Grant RGPIN-2017-03804), the Canada Research Chairs program, and an Early Researcher Award (ER18-14-158).

## Supplementary materials

At the end of the document

## Supplementary materials

### S1. Comparison between load effects during encoding and maintenance

Stronger positive load-modulated connectivity during the maintenance phase compared to the encoding phase can at least party be explained by the longer duration of the interval. When splitting the maintenance interval in 4 periods of 2 seconds (i.e., the same duration as the encoding phase), we see comparable connectivity strength during each of the 4 maintenance windows as during encoding. Figure S1 shows the unthresholded matrices for load-modulated connectivity (ss3 > ss1) and Figure S2 the average per network pair. Thus, the apparent difference in connectivity strength between encoding and maintenance may simply be an artefact of the difference in timing between these two phases.

**Figure S1.1.**
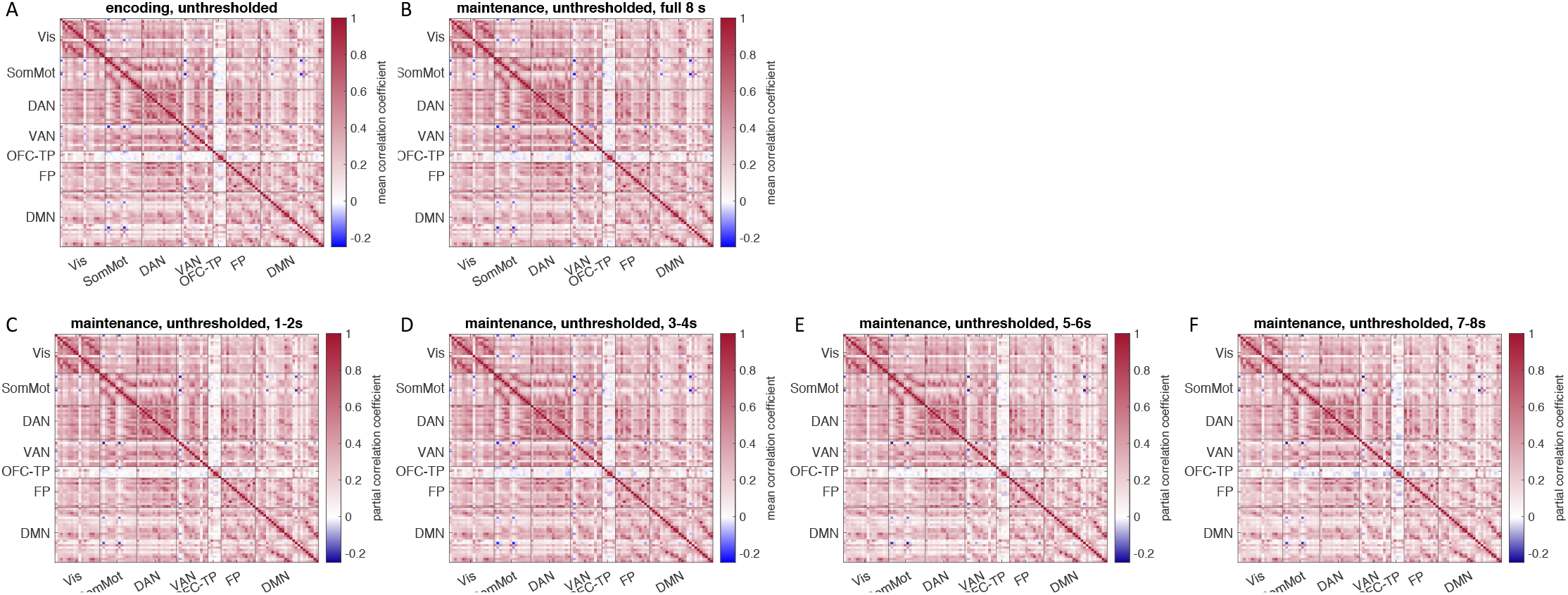
Partial correlation coefficient of load-modulated connectivity (ss3 > ss1), unthresholded. Panel A) encoding, and B) maintenance during the full 8 seconds. Panels C-F) maintenance split up in 2 second windows.

**Figure S1.2.**
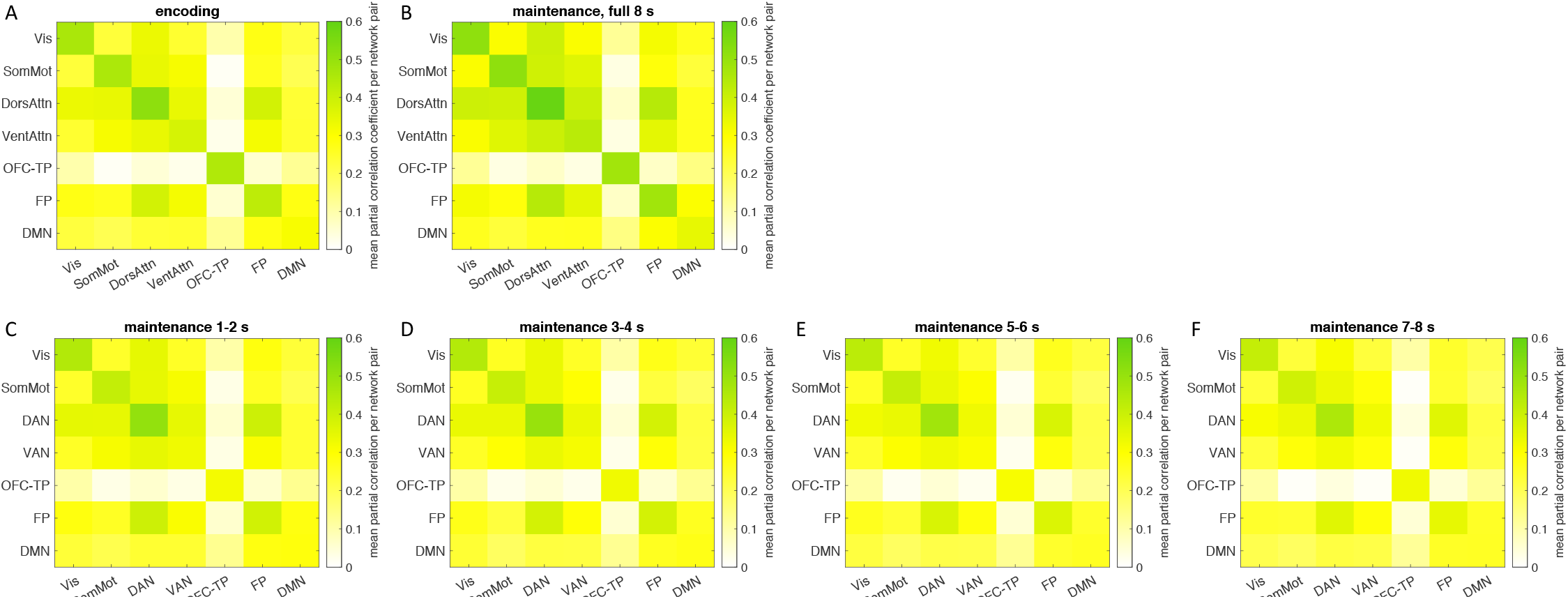
Partial correlation coefficient of load-modulated connectivity (ss3 > ss1), unthresholded, average per network pair. Panel A) encoding, and B) maintenance during the full 8 seconds. Panels C-F) maintenance split up in 2 second windows.

### S2. Supplementary analyses of the effect of performance on load-modulated functional connectivity

When only testing the effect of performance within the visual and dorsal attention networks, we found a negative relationship between response error (higher response error indicating worse performance) and connectivity within the dorsal attention network. Figure S3 shows this relationship for the average connectivity within the dorsal attention network.

The effect of memory load was most pronounced within the visual and dorsal attention networks, with only connectivity in the dorsal attention network relating to performance. However, other within and between network connections also survived thresholding for the effect of load (Figure 3 B and E). In a further, exploratory analysis, we therefore tested whether connectivity is associated with performance when including all surviving edges for the effect of load (Figure 3 B and E). This analysis replicated our previous results, showing most edges related to performance within the dorsal attention network and a few within the frontoparietal network, and between these networks (Figure S2.2). However, the results did not survive correction for testing positive and negative effects separately (*p*<.025; encoding *p* = .031 maintenance *p* = .033). The number of edges is different in both analyses (241 within the visual and dorsal attention network versus 474 and 646 edges for the effect of load during encoding and maintenance, respectively).

**Figure S2.1.**
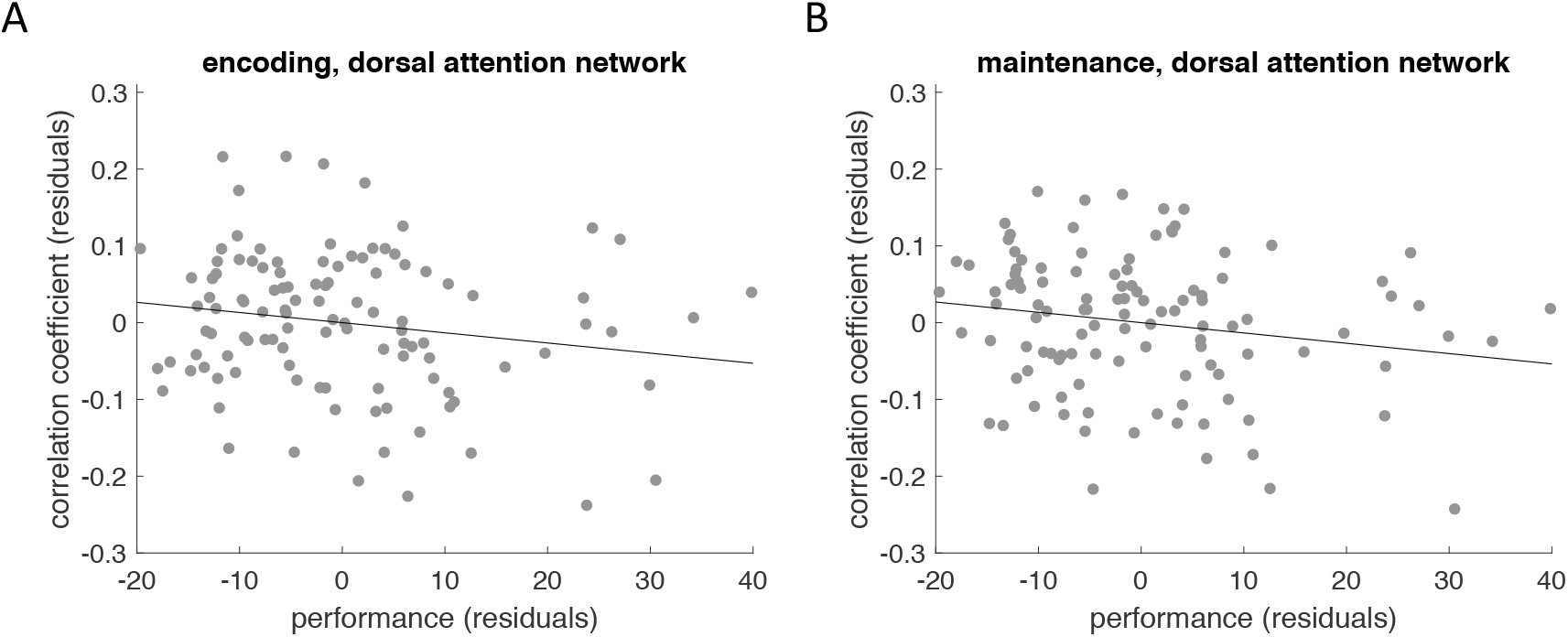
Relation between performance (response error at set size 3) and the average load-modulated connectivity strength (partial correlation coefficient) within the dorsal attention network, controlled for age. Left during encoding, right during maintenance.

**Figure S2.2.**
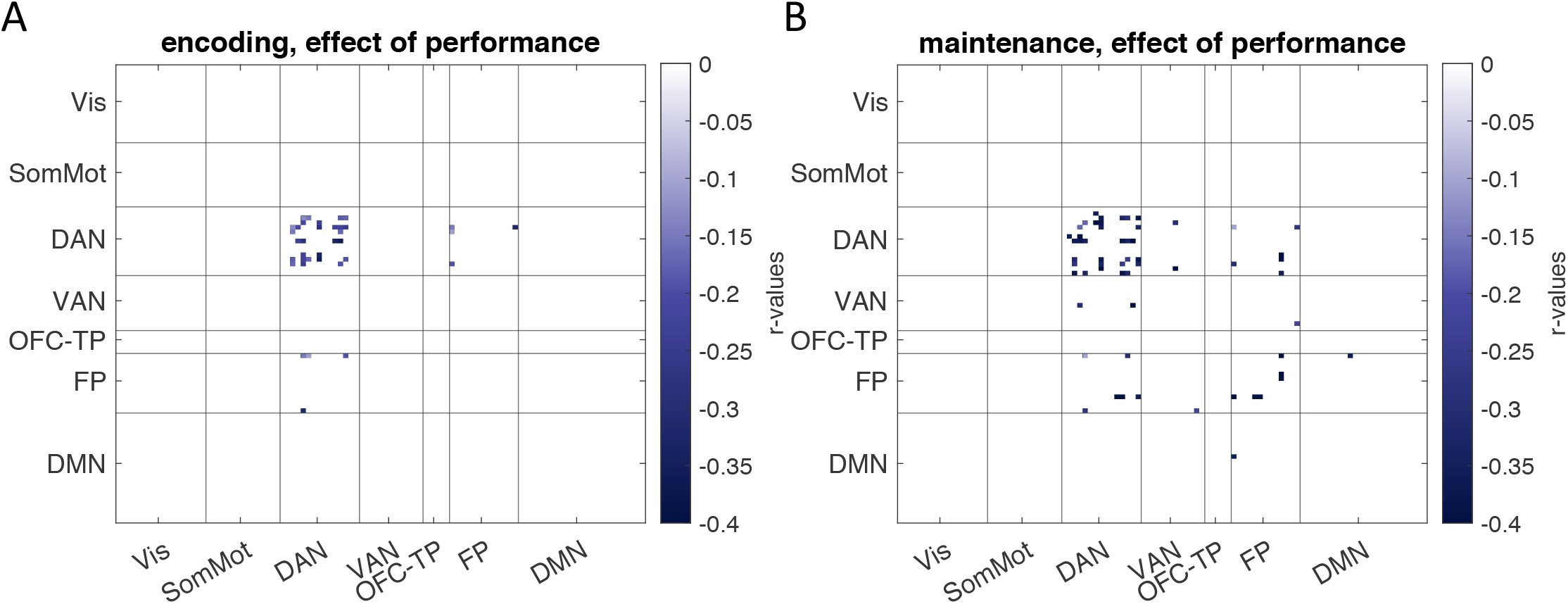
Correlation matrices for encoding (A) and maintenance (B) for the correlation between performance (response error at set size 3) and load-modulated connectivity strength (partial correlation coefficient), over and above the effect of age. Edges included in the analyses were those that showed a significant effect of load (Figure 3 B and E). These results do not survive correction for testing positive and negative effects of performance independently (encoding *p* = .031 maintenance *p* = .033).

## Footnotes

1 The effect of load may seem stronger during the maintenance period compared to the encoding period, but this appears to be due to the difference in duration between encoding and maintenance (2 versus 8 s, respectively, see supplementary materials S1).

